# Body size and food plants drive local extinction risk in butterflies

**DOI:** 10.1101/2020.08.03.233965

**Authors:** Anwar Palash, Shatabdi Paul, Sabrina Karim Resha, Md Kawsar Khan

## Abstract

Lepidoptera, butterflies and moths, are significant pollinators and ecosystem health indicators. Therefore, monitoring their diversity, distribution, and extinction risks are of critical importance. We aim to understand the drivers of the local extinction risks of the butterflies in Bangladesh. We conducted a systematic review to extract the diversity, distribution and local extinction risks of the butterflies of Bangladesh, and possible drivers of their extinction, e.g., body size, host plants and nectar plants. We updated the current checklist, which now consists of 463 species. We provided distribution and extinctions risk atlas showing both the diversity and extinction risks were highest in the eastern region of Bangladesh. We tested whether body size and host plants contribute to the local extinction risks of butterflies. We predicted butterflies with larger body size and fewer host plants and nectar plants would be in greater extinction risk. Accordingly, we showed that extinction risk was higher in larger butterflies than smaller butterflies, and in butterflies with a fewer number of host plants and nectar plants than the butterflies with higher number host plants. Our study highlights the contribution of body size and host plants as potential drivers of the local extinction risks of butterflies.

## Introduction

Biodiversity— the variety of life, including variation among genes, species, and functional traits— is crucial for the survival of the whole sphere of life on earth [1,2]. Despite continuous conservation efforts, most indicators of the state of biodiversity (covering species’ population trends, extinction risk, habitat extent and condition, and community composition) are on the decline. On the other hand, indicators of pressures on biodiversity (including resource consumption, invasive alien species, nitrogen pollution, overexploitation, and climate change impacts) are on the rise [1] resulting biodiversity loss which might affect the dynamics and functioning of ecosystems [3]. Sustaining biodiversity is a fundamental process to maintain a functional ecosystem. Continuous monitoring of extinction risks and understanding the roles its drivers are essential to conserve species and to maintain ecological balance.

The regional distribution of a species is thought to be a combined outcome of colonization and extinction of a set of local populations [4]. Extinction of species and the reduction of species’ distribution areas are primarily caused by anthropogenic activities and because of climate change [5]. Recent studies have shown nonrandom extinctions in disturbed landscapes across the world [6–8], which implies that some species may be more prone to extinction than others inherently [9]. However, we know enough to conclude that there is a lack of data on species distributions, abundance and associated ecological factors which strongly affects key biodiversity statistics including determining their extinction risk [10].

Butterflies have been a significant focus for ecologists because of their role as pollinators and ecosystem health indicators [11,12]. Butterflies are a widely distributed insect order that evolved from the early Jurassic period. A total of 180,000 species are distributed worldwide excluding Antarctica [13]. They are found in all global territory and populate in desert land even in tropical forests too [14]. However, many species of this insect group are in risk of extinction because of habitat loss and under the influence of changing climatic condition [15]. Extinction risks of butterflies have been conducted in regional scales as well global scale. Assessing the regional extinction risks and its contributing factors can assist in protecting butterflies in a particular ecogeographic region. Physiological conditions (body size, sexual dimorphism), behaviour (migration), ecological factors (host plants and nectar plants) are some factors that contribute to the extinction risks of butterflies [9]. However, studies resulted in conflicting outcomes: some studies suggested a correlation of body size, the number of host and nectar plants with extinction risks whereas other studies indicated otherwise [9,16–19], indicating the demand of further studies.

Bangladesh, despite its small geographical area, has a rich floral and faunal diversity, especially insect diversity [20]. Larsen (2004) reported the occurrence of 430 butterflies from Bangladesh and predicted the number of species should be 500-550 [21]. Since then, several species were added to Bangladeshi butterfly fauna [22,23], although the checklist has not been updated. There had been vast unconsolidated data on Bangladeshi butterflies that are needed to be compiled in a digitized form so that the data could be implemented in evolution, ecology, taxonomy, and conservation research. Furthermore, distribution, life history, and associated ecological factors could be used to understand the extinction risks of different species and to protect endangered species and their habitats.

Here, we aim to annotate the diversity, distribution, and extinction risks of the Bangladeshi butterflies. We systematically search for the publications focusing on the butterflies in Bangladesh and amassed occurrence records from published articles. We updated the checklist of the butterflies of Bangladesh and provided a distribution atlas based on those records. Next, we extracted local extinction risks of the butterflies of Bangladesh from IUCN red list and provided a distribution atlas of the endangered butterflies. We then extracted data body size, larval host plants, and adult nectar plants of the butterflies to understand the contributions of these factors to local extinct risks. We predicted that extinction risks of the butterflies with larger body size, fewer host plants, and fewer nectar plants would be greater than that of butterflies with smaller body size, higher host plants, and nectar plants.

## Methods

### Ethical statement

We collected data from published articles based on our systematic literature search. We did not require ethics approval to conduct the study because we did not collect data on any endangered species and did not conduct fieldwork in national parks or protected areas.

### Literature review

We conducted a systematic literature search in the Web of Science database to identify studies on the butterflies of Bangladesh. The search was conducted on December 01, 2019, using the keywords “butterfly” and “Bangladesh” and articles published since 1971 were extracted. The search was later updated on May 31, 2020, and then the updated findings were merged with the initial search. We further cross-checked retrieved articles to identify further relevant articles that we might have missed during the Web of Science search.

### Study selection criteria

We included publications for this study if 1) studies were conducted inside the geographical distribution of Bangladesh 2) articles reported body size data of butterflies 3) publications were focused on diversity and distribution of butterflies, 4) publications focused on host plants of the larval stage of butterflies and food sources of adult butterflies, 5) articles, written in English, 6) studies published between 1971 to 2020, and 7) publications available in full-text. We retrieved the full-text of the articles and selected articles based on the inclusion criteria. Furthermore, we also used full-text for cross-referencing and extracted relevant articles.

### Checklist, geographic coverage, and occurrence data

We compiled an updated checklist of the butterflies of Bangladesh by incorporating all the species recorded within the geographical distribution of Bangladesh. We updated the taxonomic status, classifications, and synonyms when applicable by following Smetacek (2017).

We classified the geographical distribution of the butterflies of Bangladesh into three levels according to their administrative regions (i.e. division, district, and coordinates level) (Shah & Khan, 2020). We first extracted division and district level occurrence data on the Bangladeshi Lepidoptera from the published articles. We further extracted finer level occurrence data by extracting coordinates of the occurrence of butterflies. We extracted species-specific geographical coordinates when they were provided in the publications. Otherwise, we used Google maps (www.google.com/maps) to determine the longitude and latitude of a location [25]. When administrative districts were reported as locations of occurrence, we extracted longitude and latitude of the centre of the region [26].

### Wing size

We extracted the wingspan of the butterflies from the published sources (S1). We took the middle value when wing size was provided in a range. All values were recorded in millimeters (mm). Values were converted to the millimeter when they were presented in other units.

### Host plants, nectar plants, and extinction risk

We extracted data for butterflies’ host plants — plants in which, butterflies lay eggs and butterflies larva complete their life cycle. We also extracted data on nectar plants – plants from which adult butterflies collect nectar. We collected the threat status of the butterflies of Bangladesh from the IUCN Red List of Bangladesh (2015). The IUCN Red List of Threatened Species categorizes extinction risk with the following categories: least concern, near threatened, vulnerable, and critically endangered. We converted the risk categories to a numeric index from least concern (1) to critically endangered (5).

### Statistical analysis

To determine if the body size of butterflies was correlated with regional extinction risk, we applied a mixed-effect model with extinction risk as a response variable and body size as a covariate and butterfly families as a random factor. Similarly, we applied a mixed-effect model with extinction risk as a response variable and the number of nectar plants as covariate and butterfly families as a random factor. R square values of the mixed effect models were determined by using r.squareGLMM function of MuMIn package [28]. All the data were analyzed in RStudio ver 3.6.2 (R Core Team, 2019) using packages lme4 [30], dplyr [31].

## Results

### Study selection

The systematic search returned a total of 92 publications (Fig 1). After an initial review of the titles and abstracts, 18 publications were excluded because they did not meet the selection criteria (studies did not focus on butterflies or conducted outside of the geographical distribution of Bangladesh) (Fig 1). The full text of the 74 publications was obtained and screened for the selection criteria (Fig 1). Of these, 18 publications were excluded because those studies did not provide data for the distribution, wing size, host plants, and food plants of the butterflies of Bangladesh (Fig 1). Finally, 56 publications fulfilled the selection criteria (Fig 1). A full list of all publications included in this study is provided in the supplementary data (Supplementary file S1).

**Fig 1.**
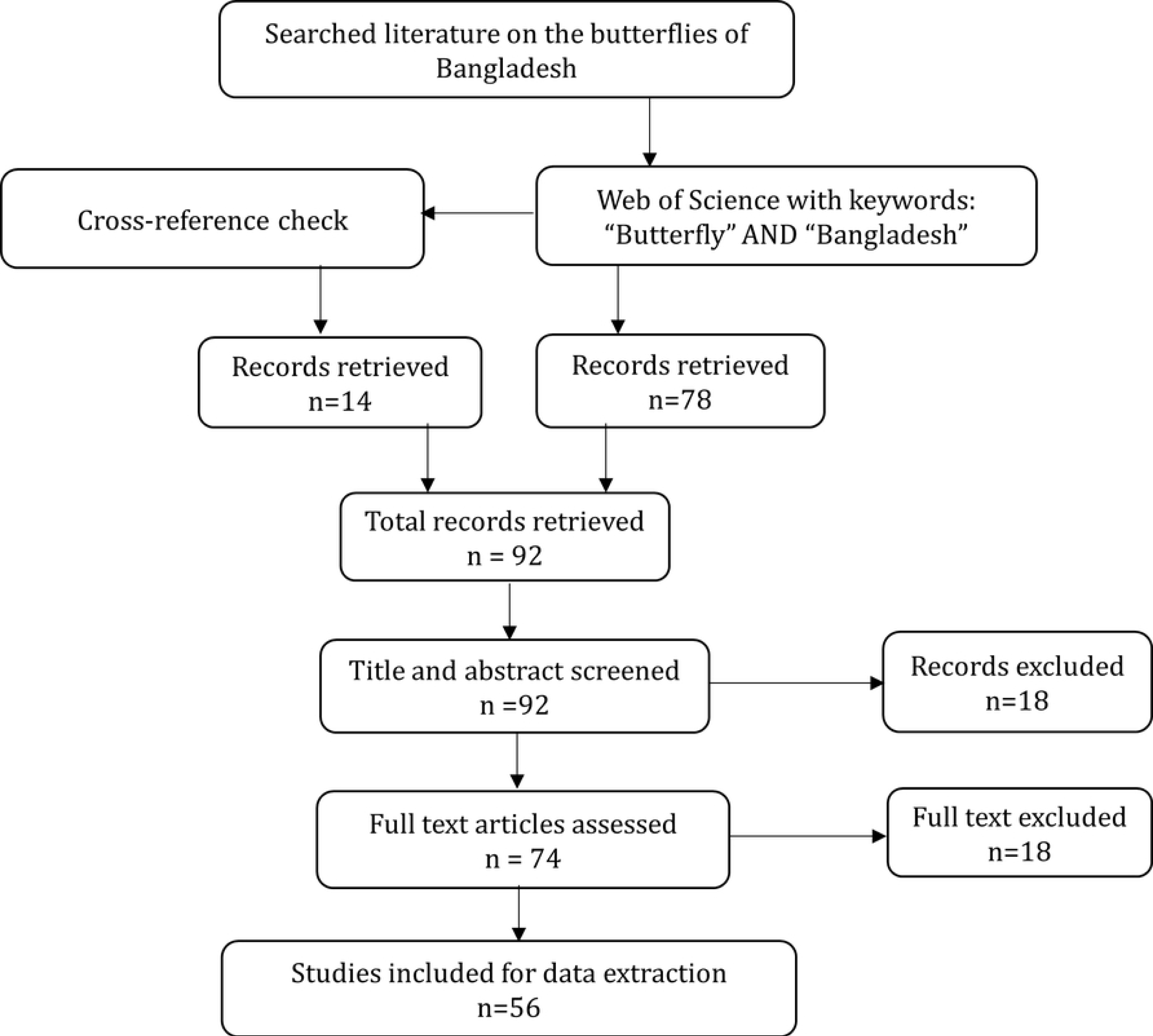
Schematic overview of the systematic literature search.

### Data extraction

Our systematic review showed the occurrence of 463 butterflies in Bangladesh. We extracted the wingspan size of 268 species (Supplementary file S2). We also recorded larval host plants of 183 species and food plants of 126 species (Supplementary file S2). Furthermore, we retrieved the local extinction risk status of 315 species (Supplementary file S2). Finally, we amassed 3622 occurrence records from 93 different locations and 44 administrative districts from 1844 to March 2020 (Supplementary file S3).

### Checklist and taxonomic coverage

We updated the checklist of the butterflies of Bangladesh, which showed the occurrence of 463 butterflies species belonging to six families. Nymphalidae was the largest family with 155 species followed by Lycaenidae family with 138 species. The rest of the four families were Hesperiidae, Pieridae, Papilionidae, Riodinidae with 93, 40, 32, and 5 species respectively.

### Distribution and conservation status

Sylhet and Chittagong were two most diverse regions with 321 and 299 recorded species whereas Barisal had the lowest species richness with 64 species (Fig 2). The central region Dhaka held a moderate number of butterflies with 216 species, despite being the most urbanized region of the country (Fig 2).

**Fig 2.**
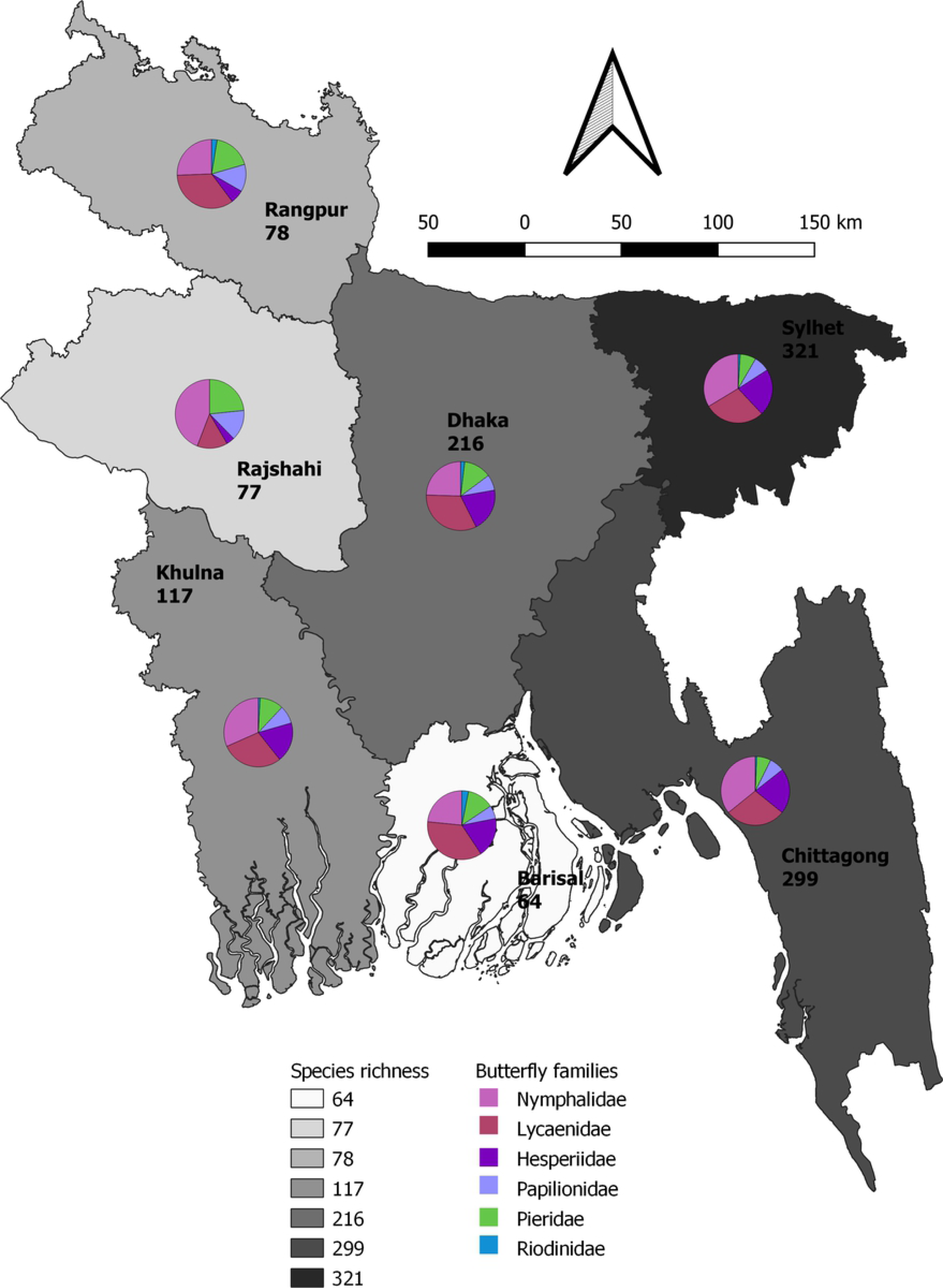
A reference map showing butterfly species richness in seven divisions in Bangladesh. The background colour of each division represent species richness of that region. Pie charts illustrate the proportional composition of butterfly families in each division.

The national IUCN Red List status showed one species (*Euploea crameri* Moore, 1890) as Critically Endangered (CR) and 112 species as Endangered (EN). The number of Vulnerable (VU), Least Concern (LC), and Data Deficient (DD) species were 81, 94, and 27 respectively.

The highest number of threatened species (CR, EN, VU) were recorded in Sylhet division (151 species) followed by Chittagong (137 species) (Fig 3). Almost 40% and 36% of the species found in Dhaka (86) and Khulna (42) falls under the Threatened category (Fig 3).

**Fig 3.**
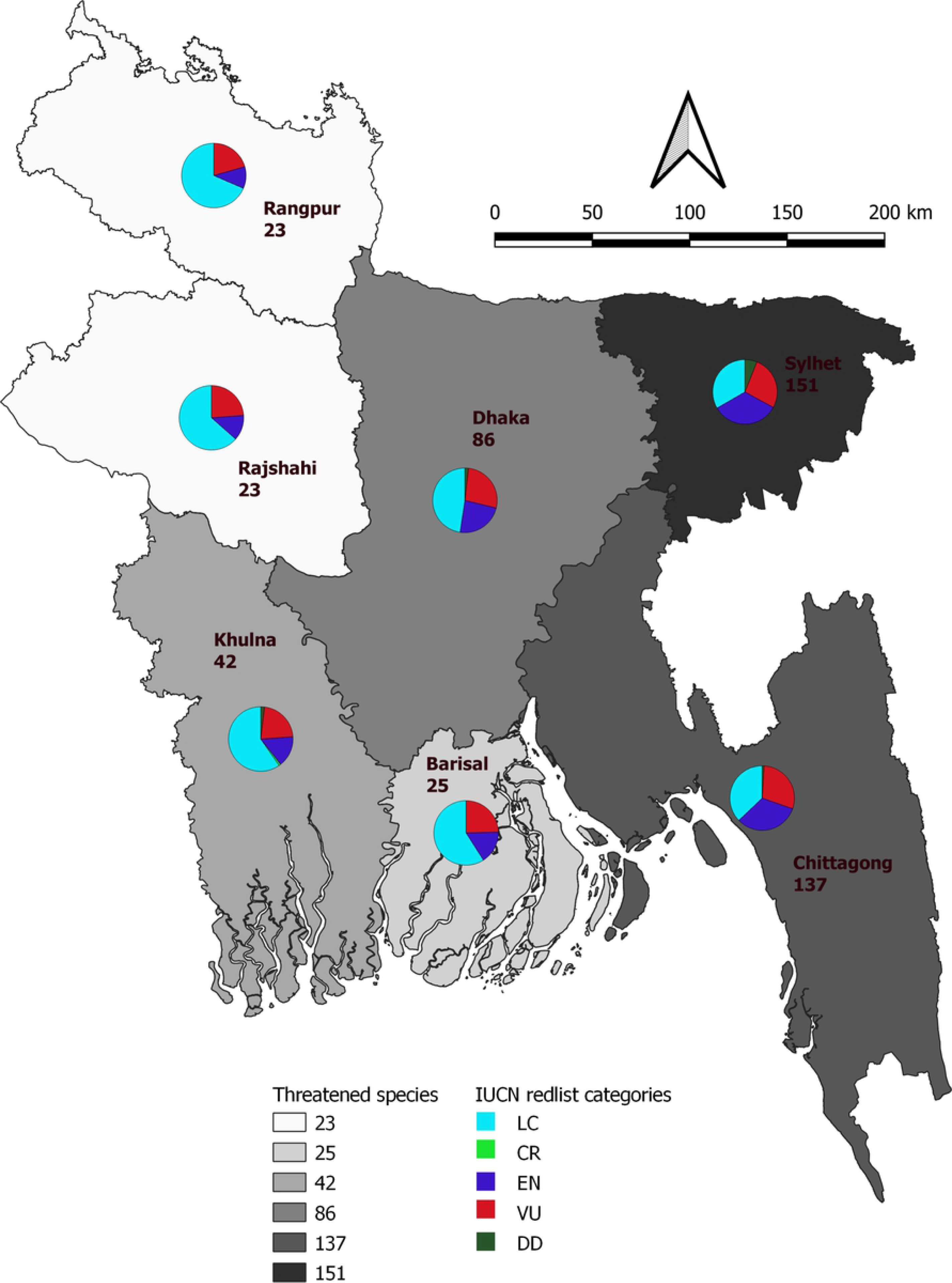
A reference map showing butterfly species richness of the threatened category butterflies in seven divisions in Bangladesh. The background colour of each division represent species richness of that region. Pie charts illustrate the proportional composition of butterfly families in each division.

### Threat status

The extinction risk of the butterflies was correlated with their wingspan; larger butterflies were in higher extinction risk than smaller butterflies (LME: estimate = 1.5286 ± 0.7284, df = 233.52, t =2.09, p = 0.036, R^2^ = 0.72; Fig 4a). Higher number of host plant reduced extinction risks of butterflies; butterflies with higher number of host plants were in lower extinction risks than butterflies with lower host plants (LME: estimate = −0.17201 ± 0.02922, df = 179, t =-5.88, p < 0.0001, R^2^ = 0.15; Fig 4b). Similarly, butterflies feeding on larger number of nectar plants were in lower extinction risk compared to butterflies feeding on fewer number of nectar plants (LME: estimate = −0.16405 ± 0.04704, df = 114.60, t =-3.48, p <0.001, R^2^ = 0.13, Fig 4c).

**Fig 4:**
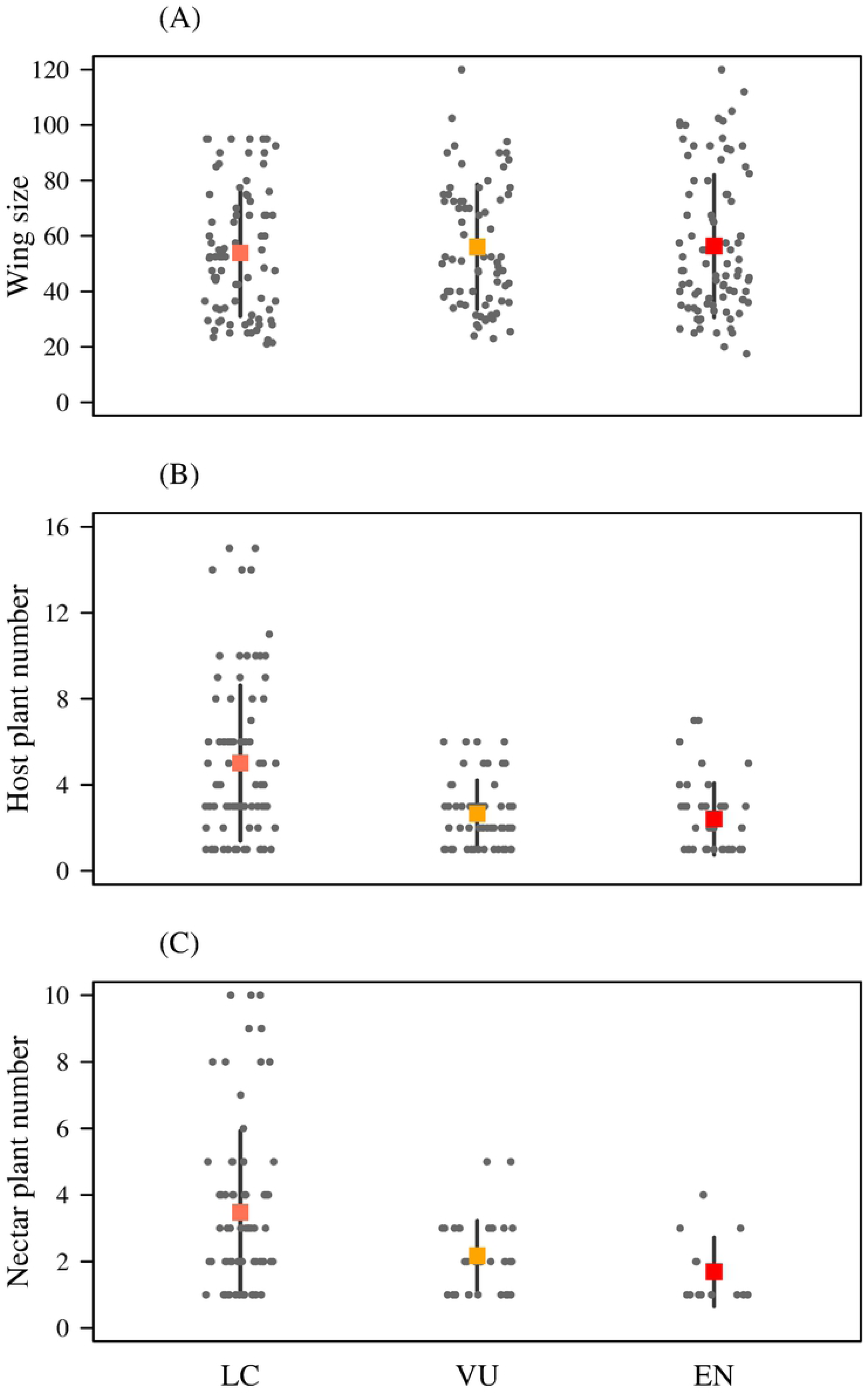
Contributions of body size and food plant to local extinction risk of butterflies. A) wing size (mean ± sd) and local extinction risks of butterflies, B) host plant number (mean ± sd) and local extinction risk of butterflies, and C) nectar plant number (mean ± sd) and local extinction risk of butterflies. Squares indicate mean of data; horizontal lines indicate standard deviation (sd) of the data; circles indicate individuals data point. LC= least concern, VU = vulnerable, and EN= endangered.

## Discussion

Monitoring diversity, abundance, and extinction risks of butterflies in regional scales are essential because of their contributions as pollinators. We updated the checklist of the butterflies of Bangladesh and extracted their distribution records from previously published articles. We showed the diversity of Butterflies is highest in the eastern region of Bangladesh. We tested contributions of the body size and food plants of the butterflies to their extinction risks. We found that larger butterflies are in greater extinction risk than the smaller butterflies. We further showed that extinction risks of butterflies with fewer host and nectar plants are greater than butterflies with higher host and nectar plants.

Bangladesh has high Butterflies diversity, despite being a small country. Earlier studies by Larsen, (2004) confirmed the presence of 427 Butterflies species in Bangladesh and suggested that the total number of species could be between 500 – 550. Since then, the checklist was not updated, although several articles recorded new species from Bangladesh [22,23]. Our updated list showed the number of Butterflies species in Bangladesh reached 463, which has increased by 36 species since 2004. These new additions indicate the importance of continuous biodiversity assessment and suggest that the number of Butterflies in Bangladesh might increase with more studies.

Monitoring temporal and spatial distribution is essential for the conservation of species, especially for species with a short life cycle, small geographical distribution, and a strict habitat requirement such as Butterflies [26]. We showed that the eastern regions of Bangladesh had greater Butterflies diversity. Bangladesh is situated at the intersection of the Indo-Himalayan and Indo-Chinese sub-region, in the transitional zone for flora and fauna of the Indian subcontinent and Southeast Asia, and the eastern region is part of the Indo-Burma biodiversity hotspot contains greater habitat diversity compared to the other parts of Bangladesh [32–34]. These habitat diversities, including the presence of deciduous tropical forests, semi-evergreen, and evergreen forests, with fragmented forest patches, provide suitable habitats for many butterflies [21]. Furthermore, the high plant diversity in this region provides a greater source of host plants and nectar plants for many species thereby contributing to the greater butterfly diversity [21]. Similarly, a higher diversity of other invertebrates such as Odonata, and Diptera were also recorded from this region [35–37]. We found lower butterfly diversity in Barisal, Khulna, Rajshahi, and Rangpur compared to other parts of Bangladesh. These regions are also studied less compared to other parts thereby suggesting lower diversity could occur because of less habitat diversity or due to fewer studies. Further studies are required to pinpoint the underlying causes.

Understanding the extinction risk of a species is essential to develop conservation strategies for the protection of a species. Animal body size is correlated with several species attributes such as longevity, reproduction, resource use, and average population density [17]; all of which contribute to the extinction risk. Analysis of invertebrates [38–41] and vertebrates [42,43] body size on global and local scale suggests that larger species are usually more vulnerable to extinction. Previous studies on the impact of Lepidopteran wingspan suggest that butterfly wingspan is neither significant [9], nor primary [44] determinant of extinction. Furthermore, Nieminen (1996) concluded that moth size did not affect the risk of population extinction independently. We showed the larger Bangladeshi butterflies are in greater local extinction risk but it was not statistically significant. Our finding corroborates previous studies.

The diversity of butterflies, like many other insects, depends on the abundance of food sources and habitats [45,46]. Butterfly larvae feed on host plant leaves and adult butterflies collect nectar from multiple plant sources, therefore higher plant richness promotes greater butterfly diversity [47]. Previous studies showed that larval host plant dynamics are the most important determinant regional extinction of butterflies [9,48]. Similarly, we found that larval host plant dynamics to be the most significant determinant of butterflies’ extinction. We found butterflies with fewer host plants have higher extinction risk. Highly host-specific butterflies are more dependent on the survival of host plants than less host-specific butterflies and are more likely to be vulnerable to localized fragmentation of resources [9]. Furthermore, co-extinction of butterflies are likely to occur with their host plants [19,49].

The second most significant factor that determined the extinction risk of butterflies was nectar plants partly because of the same reason as the host plants since many of the host plants act as the nectar plants and food source for many butterflies [50]. Butterflies depend on nectar sources to mature their eggs [11]. Besides, adult nectar feeding of butterflies has long been shown to be critically important for somatic maintenance and reproduction, and population persistence [51,52], especially for the species that emerge with unyoked eggs [53]. Moreover, seasonal variation in nectar availability may induce local extinction [11]. As a result, the number of nectar plants and food resources is particularly essential for the survival of butterflies. So, our finding of butterflies feeding on a larger number of nectar plants are in lower extinction risk compared to butterflies feeding on a fewer number of nectar plants strengthens the previous findings. There is a comparative lack of study of nectar plants of Bangladeshi butterflies than the host plant. With a similar effort of research, the significance of nectar plants would probably be closely similar to host plants.

Understanding the spatial distribution and monitoring regional species diversity regularly is essential to understand the population trends, extinction risks of insects. In this study, we extracted occurrence and distribution data on the butterflies of Bangladesh from published literature, field guides, and our unpublished data. We updated the checklist and atlas of the butterflies of Bangladesh. We analyzed the regional extinction risk of the butterflies and found that butterflies with larger wingspan, fewer host plants, and nectar plants are in greater extinction risk. Our study highlights vulnerable butterflies in Bangladesh and the drivers associated with their extinction risks. Our study will provide resources to develop conservation strategies to protect the butterflies in Bangladesh and beyond.

## Acknowledgement

MKK thanks Payal Barua and Leelaboti Khona for their support. Conflict of interest: The authors declare no conflict of interest.

## Data accessibility

All data will be uploaded in figshare depository upon acceptance of the paper.

## References

1. Butchart SH, Walpole M, Collen B, Van Strien A, Scharlemann JP, Almond RE, et al. Global biodiversity: indicators of recent declines. Science. 2010;328: 1164–1168.

2. Gonçalves J, Alves P, Pôças I, Marcos B, Sousa-Silva R, Lomba Â, et al. Exploring the spatiotemporal dynamics of habitat suitability to improve conservation management of a vulnerable plant species. Biodivers Conserv. 2016;25: 2867–2888.

3. Cardinale BJ, Duffy JE, Gonzalez A, Hooper DU, Perrings C, Venail P, et al. Biodiversity loss and its impact on humanity. Nature. 2012;486: 59–67.

4. Levins R. Some demographic and genetic consequences of environmental heterogeneity for biological control. Am Entomol. 1969;15: 237–240.

5. Tsahar E, Izhaki I, Lev-Yadun S, Bar-Oz G. Distribution and extinction of ungulates during the Holocene of the southern Levant. PLoS One. 2009;4: e5316.

6. McKinney ML. Extinction vulnerability and selectivity: combining ecological and paleontological views. Annu Rev Ecol Syst. 1997;28: 495–516.

7. Russell GJ, Brooks TM, McKinney MM, Anderson CG. Present and future taxonomic selectivity in bird and mammal extinctions. Conserv Biol. 1998;12: 1365–1376.

8. Purvis A, Gittleman JL, Cowlishaw G, Mace GM. Predicting extinction risk in declining species. Proc R Soc Lond B Biol Sci. 2000;267: 1947–1952.

9. Koh LP, Sodhi NS, Brook BW. Ecological correlates of extinction proneness in tropical butterflies. Conserv Biol. 2004;18: 1571–1578.

10. Pimm SL, Jenkins CN, Abell R, Brooks TM, Gittleman JL, Joppa LN, et al. The biodiversity of species and their rates of extinction, distribution, and protection. science. 2014;344.

11. New TR. Lepidoptera and conservation. John Wiley & Sons; 2013.

12. Proctor M, Yeo P, Lack A. The natural history of pollination. HarperCollins Publishers; 1996.

13. Capinera JL. Encyclopedia of entomology. Springer Science & Business Media; 2008.

14. Gullan PJ, Cranston PS. The insects: an outline of entomology. John Wiley & Sons; 2014.

15. Franzén M, Johannesson M. Predicting extinction risk of butterflies and moths (Macrolepidoptera) from distribution patterns and species characteristics. J Insect Conserv. 2007;11: 367–390. doi: 10.1007/s10841-006-9053-6

16. Nieminen M. Risk of population extinction in moths: effect of host plant characteristics. Oikos. 1996; 475–484.

17. Johst K, Brandl R. Body size and extinction risk in a stochastic environment. Oikos. 1997; 612–617.

18. Hanski I, Singer MC. Extinction-colonization dynamics and host-plant choice in butterfly metapopulations. Am Nat. 2001;158: 341–353.

19. Koh LP, Sodhi NS, Brook BW. Co-extinctions of tropical butterflies and their hostplants. Biotropica. 2004;36: 272–274.

20. Shah MNA, Khan MK. OdoBD: An online database for the dragonflies and damselflies of Bangladesh. PloS One. 2020;15: e0231727. doi: https://doi.org/10.1371/journal.pone.0231727

21. Larsen TB. Butterflies of Bangladesh: an annotated checklist. IUCN, the World Conservation Union, Bangladesh Country Office; 2004.

22. Khan AKMMA, Khan T, Khan MK. Three new records of Butterfly from north-east region of Bangladesh. Festschr 50th Anniv IUCN Red List Threat Species. 2014; 35–38.

23. Khan K. Three new records of butterfly from university of Chittagong and Shahjalal University of science and technology in Bangladesh. Int J Fauna Biol Stud. 2014;1: 30–33.

24. Smetacek P. A Naturalist’s Guide to the Butterflies of India, Pakistan, Nepal, Bhutan, Bangladesh and Sri Lanka. John Beaufoy Publishing Limited; 2017.

25. Zhang G, Zheng D, Tian Y, Li S. A dataset of distribution and diversity of ticks in China. Sci Data. 2019;6: 1–7.

26. Shah MNA, Khan MK. OdoBD: An online database for the dragonflies and damselflies of Bangladesh. Plos One. 2020;15: e0231727.

27. Red List of Bangladesh : volume 7 : butterflies. In: IUCN [Internet]. 7 Sep 2016 [cited 22 May 2020]. Available: https://www.iucn.org/content/red-list-bangladesh-volume-7-butterflies

28. Barton K, Barton MK. Package ‘MuMIn.’ R Package Version. 2019;1.

29. Team RC. R: A language and environment for statistical computing. 2013.

30. Bates D, Sarkar D, Bates MD, Matrix L. The lme4 package. R Package Version. 2007;2: 74.

31. Wickham H, François R, Henry L, Müller K, RStudio. dplyr: A Grammar of Data Manipulation. 2019. Available: https://CRAN.R-project.org/package=dplyr

32. Stanford CB. The capped langur in Bangladesh: behavioral ecology and reproductive tactics. Karger Medical and Scientific Publishers; 1991.

33. Tordoff AW, Bezuijen MR, Duckworth JW, Fellowes JR, Koenig K, Pollard EHB, et al. Ecosystem profile: Indo-Burma biodiversity hotspot, 2011 update. Crit Ecosyst Partnersh Fund Conserv Int Arlingt Xxi 360pp. 2012.

34. Feeroz MM. Biodiversity of Protected Areas of Bangladesh, Vol. III: Teknaf Wildlife Sanctuary. Bio Track Arannayk Found Dhaka 240pp. 2013.

35. Khan MK. Dragonflies and damselflies (Insecta: Odonata) of the northeastern region of Bangladesh with five new additions to the Odonata fauna of Bangladesh. J Threat Taxa. 2015;7: 7795–7804. doi: 10.11609/JoTT.o4314.7795-804

36. Khan MK. Odonata of eastern Bangladesh with three new records for the country. J Threat Taxa. 2018;10: 12821–12827. doi: 10.11609/jott.3819.10.13.12821-12827

37. Leblanc L, Hossain MA, Doorenweerd C, Khan SA, Momen M, Jose MS, et al. Six years of fruit fly surveys in Bangladesh: a new species, 33 new country records and discovery of the highly invasive Bactrocera carambolae (Diptera, Tephritidae). ZooKeys. 2019;876: 87–109. doi: 10.3897/zookeys.876.38096

38. Mattila N, Kotiaho JS, Kaitala V, Komonen A. The use of ecological traits in extinction risk assessments: a case study on geometrid moths. Biol Conserv. 2008;141: 2322–2328.

39. Seibold S, Brandl R, Buse J, Hothorn T, Schmidl J, Thorn S, et al. Association of extinction risk of saproxylic beetles with ecological degradation of forests in Europe. Conserv Biol. 2015;29: 382–390.

40. Shahabuddin G, Ponte CA. Frugivorous butterfly species in tropical forest fragments: correlates of vulnerability to extinction. Biodivers Conserv. 2005;14: 1137–1152.

41. Terzopoulou S, Rigal F, Whittaker RJ, Borges PA, Triantis KA. Drivers of extinction: the case of Azorean beetles. Biol Lett. 2015;11: 20150273.

42. Bennett PM, Owens IP. Variation in extinction risk among birds: chance or evolutionary predisposition? Proc R Soc Lond B Biol Sci. 1997;264: 401–408.

43. Fritz SA, Bininda-Emonds OR, Purvis A. Geographical variation in predictors of mammalian extinction risk: big is bad, but only in the tropics. Ecol Lett. 2009;12: 538–549.

44. Kingsolver JG. Experimental analyses of wing size, flight, and survival in the western white butterfly. Evolution. 1999;53: 1479–1490.

45. Burghardt KT, Tallamy DW, Gregory Shriver W. Impact of native plants on bird and butterfly biodiversity in suburban landscapes. Conserv Biol. 2009;23: 219–224.

46. Ricketts TH, Daily GC, Ehrlich PR. Does butterfly diversity predict moth diversity? Testing a popular indicator taxon at local scales. Biol Conserv. 2002;103: 361–370.

47. Bashar MA, Mamun MA, Aslam AFM, Chowdhury AK. Biodiversity maintenance and conservation of butterfly-plant association in some forests of Bangladesh. Bangladesh J Zool. 2006;34: 55.

48. León-Cortés JL, Lennon JJ, Thomas CD. Ecological dynamics of extinct species in empty habitat networks. 2. The role of host plant dynamics. Oikos. 2003;102: 465–477.

49. Stork NE, Lyal CH. Extinction or’co-extinction’rates? Nature. 1993;366: 307–307.

50. Dennis RLH. Just how important are structural elements as habitat components? Indications from a declining lycaenid butterfly with priority conservation status. J Insect Conserv. 2004;8: 37–45. doi: 10.1023/B:JICO.0000027496.82631.4b

51. Murphy DD, Menninger MS, Ehrlich PR. Nectar source distribution as a determinant of oviposition host species in Euphydryas chalcedona. Oecologia. 1984;62: 269–271.

52. Tudor O, Dennis RLH, Greatorex-Davies JN, Sparks TH. Flower preferences of woodland butterflies in the UK: nectaring specialists are species of conservation concern. Biol Conserv. 2004;119: 397–403.

53. Wheeler D. The role of nourishment in oogenesis. Annu Rev Entomol. 1996;41: 407–431.

